# Targeting the senescence-associated immune checkpoint GD3 ganglioside extends healthspan and blunt age-related diseases with sex-specific benefits

**DOI:** 10.64898/2026.01.06.697856

**Authors:** Iryna Moskalevska, Larisa Okorokova, Raphaël Rousset, Thierry Juhel, Eric Gilson, Bérengère Dadone-Montaudie, Laurence Bianchini, Gaël Cristofari, Julien Cherfils-Vicini

## Abstract

The accumulation and impact of senescent cells in age-related diseases are increasingly characterized. However, the mechanisms underlying their accumulation and their causative role in age-associated pathologies remain poorly understood. We recently demonstrated that senescent cells can evade immune surveillance by regulating the expression of cell surface molecules such as the disialylated ganglioside GD3, which acts as a senescence-associated immune checkpoint (SIC) ^1–3^. Targeting GD3 therefore represents a novel therapeutic opportunity for age-related diseases. Here, we examined the effects of short-term anti-GD3 antibody treatment in mid-life on aging and age-related diseases in male and female mice, revealing striking sex-specific benefits. Treatment improved healthspan, survival (+20%) and reduced non-cancer mortality in males, while in females it reduced cancer-specific mortality without significantly affecting overall survival. Anti-GD3 treatment also mitigated fibrosis in lung, liver, and kidney tissues with distinct sex-dependent responses. Importantly, these benefits persisted for over a year after treatment cessation. These findings suggest that GD3-targeted therapy holds promise as a precision approach for treating age-related diseases, with therapeutic outcomes that depend critically on biological sex.

## Main Text

### Short-term anti-GD3 immunotherapy in mid-life mice improves healthspan and modulates survival in a sex-dependent manner

Senescent cells accumulate with age and may drive many aspects of age-related pathologies and functional decline^4^. Senolytic therapies target these cells for elimination, with immunotherapies showing particular promise, including monoclonal antibodies^1,3,5^, CAR-T cells^6,7^, and peptides that restore immunosurveillance^8,9^. We recently identified the disialylated ganglioside GD3 as a senescence-associated immune checkpoint (SIC) that allows senescent cells to evade immune clearance. In our previous studies, anti-GD3 immunotherapy reduced the progression of senescence or age-related diseases, including bleomycin-induced pulmonary fibrosis^1^ and collagenase-induced bone remodeling^2^, as well as age-related bone remodeling and lung fibrosis in old male mice^1^. Here, we examined whether short-term anti-GD3 immunotherapy in mid-life could improve long-term lifespan and healthspan outcomes in both male and female mice.

As the aging process exhibits strong sexual dimorphism^10^, we evaluated the preventive effects of GD3 targeting against pathological aging in 18-month-old male and female mice. Animals were treated for 3 months with either an anti-GD3 monoclonal antibody (anti-GD3) or the reference senolytic combination Dasatinib + Quercetin (D+Q)^11,12^, administered every 15 days for a total of six doses (Fig. 1a). A subset of animals was sacrificed at 21 and 24 months for tissue and biomarker analysis, while the remaining cohort was monitored for longitudinal assessment of frailty, body weight, and survival. Frailty was evaluated using a composite score integrating clinical and physiological parameters (Extended Data Table 1). Urine and blood were collected monthly for the duration of the experiment. Overall, anti-GD3 therapy showed a trend toward increased survival (p = 0.052) and significantly improved the frailty index compared with both control and D+Q-treated mice (p < 0.001), with a 19.8% increase in 50% overall survival (OS) versus control and a 34.5% increase versus D+Q (Fig. 1b). At the median survival time, the global frailty index was reduced by 33% in anti-GD3-treated mice (Fig.1d).

**Figure 1:**
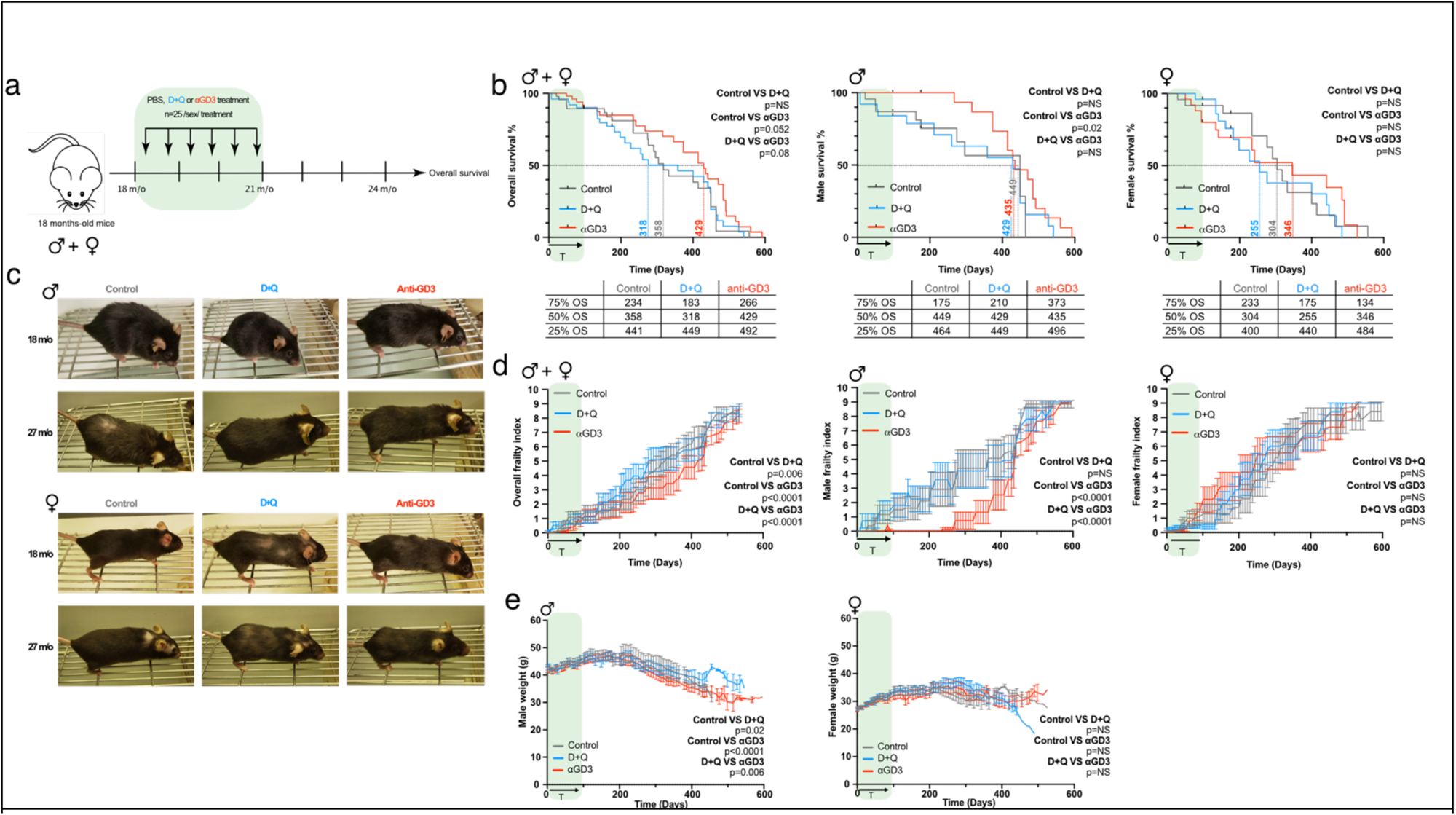
Short-term anti-GD3 immunotherapy in mid-life mice improves healthspan and modulates survival in a sex-dependent manner. **a**, Representative scheme of anti-GD3 (αGD3) or Dasatinib and Quercetin (D+Q) treatment in 18-month-old male and female mice. **b**, Overall survival (OS) analysis of all (left panel), male (middle panel) and female (right panel) mice as a function of treatment. **c**, Representative images of male (upper panel) and female (lower panel) mice at 18 and 27 months of age as a function of treatment. **d**, Evaluation of frailty index of all (left panel), male (middle panel) and female (right panel) mice in function of the treatment (+/- SEM). **e**, Weight of male (left panel) and female (right panel) mice as a function of treatment. The experiment was performed with n=25 mice per sex and per treatment. Statistical analyses: Log-rank test (b), Wilcoxon matched-pairs signed rank test (d, f).

However, sex-specific analysis revealed clear sexual dimorphism with a marked benefits in male mice, including compression of morbidity lasting over one year after treatment cessation (75% OS improved by 105% versus control) and persistent reduction in frailty for more than one year after treatment cessation (Fig. 1b, and d). In contrast, female mice displayed only modest non-significant increases in median survival (+13.8%). Anti-GD3 treatment in female mice conferred delayed survival benefits at lower percentiles (25% OS +21% versus +9.7% in males), suggesting long-lasting protection in late life despite an early drop in 75% OS (- 37%) (Fig. 1b, and d). In addition to survival and frailty, anti-GD3 therapy modulated age-related weight trajectories in a sex-dependent manner, limiting excessive weight gain in males and preventing late-life weight loss in females (Fig. 1e). Collectively, these results indicate that transient mid-life anti-GD3 therapy can durably improve healthspan and late-life survival, with sexually dimorphic effects.

### Anti-GD3 immunotherapy selectively delays non-cancer-related mortality in male mice and tumor-associated death in females without altering tumor incidence

To further dissect the impact of anti-GD3 therapy on age-related mortality, we stratified causes of death into cancer-related and non-cancer-related categories. Overall, anti-GD3 treatment predominantly influenced the median survival for non-cancer-related causes was extended by approximately 100 days in the anti-GD3 group (+39%), with no significant benefit on cancer-related deaths when pooling sexes (Fig. 2a). In male mice, anti-GD3 therapy significantly delayed non-cancer-related mortality (p=0.01 vs control), with a gain of nearly 200 days (+56%) in median survival compared to controls (Fig.2b). This delay was not associated with changes in tumor incidence or spectrum, which remained comparable across treatment arms (Fig. 2d). In female mice, the increase in median survival due to reduced non-cancer-related death was minimal (approximately 15 days), suggesting a negligible effect in this context.

**Figure 2:**
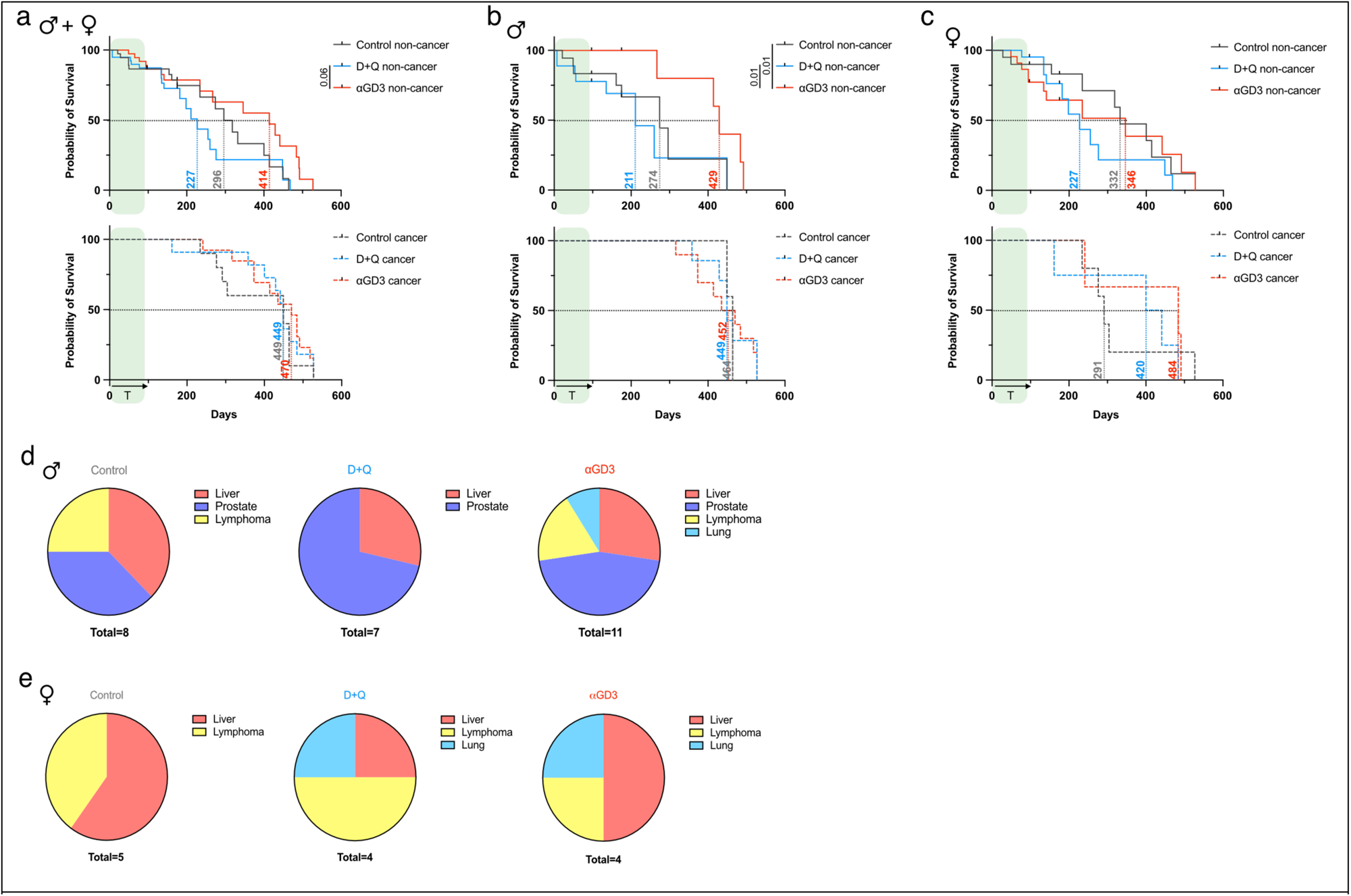
Anti-GD3 immunotherapy selectively delays non-cancer-related mortality in male mice and tumor-associated death in females without altering tumor incidence. **a-c**, Kaplan-Meier survival analysis of cause of death (cancer-induced or non-cancer-induced) and frequency of spontaneous tumor development in all (**a**), male (**b**) and female (**c**) mice as a function of treatment. Frequency of development of different cancer types in male (**d**) and female (**e**) mice in function of the treatment. The experiment was performed with n=25 mice per sex and per treatment. Statistical analysis: Log-rank test (a-c).

Conversely, analysis of cancer-related deaths revealed a sex-reversed pattern: in males, anti-GD3 had no impact on cancer-associated survival (Fig. 2b), whereas in females, both anti-GD3 and D+Q treatments markedly delayed tumor-related mortality, extending median survival by nearly 200 days (+66%) compared to controls (Fig. 2c). Despite this survival gain, tumor incidence and histopathological types (primarily involving liver, prostate, lymphoma, and lung) remained unchanged across all conditions (Fig 2d, e, Extended Data Fig. 1q).

Together, these results indicate that anti-GD3 therapy extends lifespan through distinct mechanisms in males and females: by attenuating non-cancer pathologies in males or by delaying tumor-related death in females, without altering overall tumor burden. D+Q treatment similarly delayed cancer-related deaths in females compared to anti-GD3, suggesting possible sex-specific modulation of immune surveillance mechanisms by both treatments. Importantly, blood parameters remained within the normal range throughout the treatment period in both sexes, with no signs of toxicity or systemic adverse effects (Extended Data Fig.1a–p), supporting the safety of repeated anti-GD3 administration in aged animals.

### Anti-GD3 immunotherapy attenuates age-related tissue remodeling in lung, liver and kidney in a sex-dependent manner

Given our previous data describing a role of GD3 in senescence-associated pulmonary fibrosis^1^, we first examined the effect of anti-GD3 therapy on spontaneous lung aging. Our histological analysis revealed that anti-GD3 treatment significantly reduced alveolar collagen deposition and prevented age-associated alveolar enlargement (emphysema-like changes) in both sexes (Fig. 3a–c). Notably, female mice exhibited a delayed onset of lung pathology compared to males, consistent with a less severe basal phenotype at equivalent ages. While the protective effect against emphysema was lost upon treatment cessation, the reduction in fibrotic remodeling persisted in both males and females (Fig. 3d). Mechanistically, GD3 expression levels in lung tissue decreased during treatment in both sexes, but this decrease was maintained only in male mice after treatment cessation, with GD3 expression returning to baseline in females (Fig. 3e). A similar transient dynamic was observed for Glb1, the gene coding for the well-known beta-galactosidase senescence marker, whose expression decreased during treatment but rebounded post-treatment in both sexes. We next assessed lung tumor burden. Anti-GD3 treatment reduced both the number of visible lung lesions and the proportion of proliferating (PCNA⁺) cells in both sexes (Fig. 3f–h). However, this anti-tumor effect was not sustained after therapy withdrawal. Importantly, in contrast to some D+Q regimens^13^, anti-GD3 did not promote tumor emergence.

**Figure 3:**
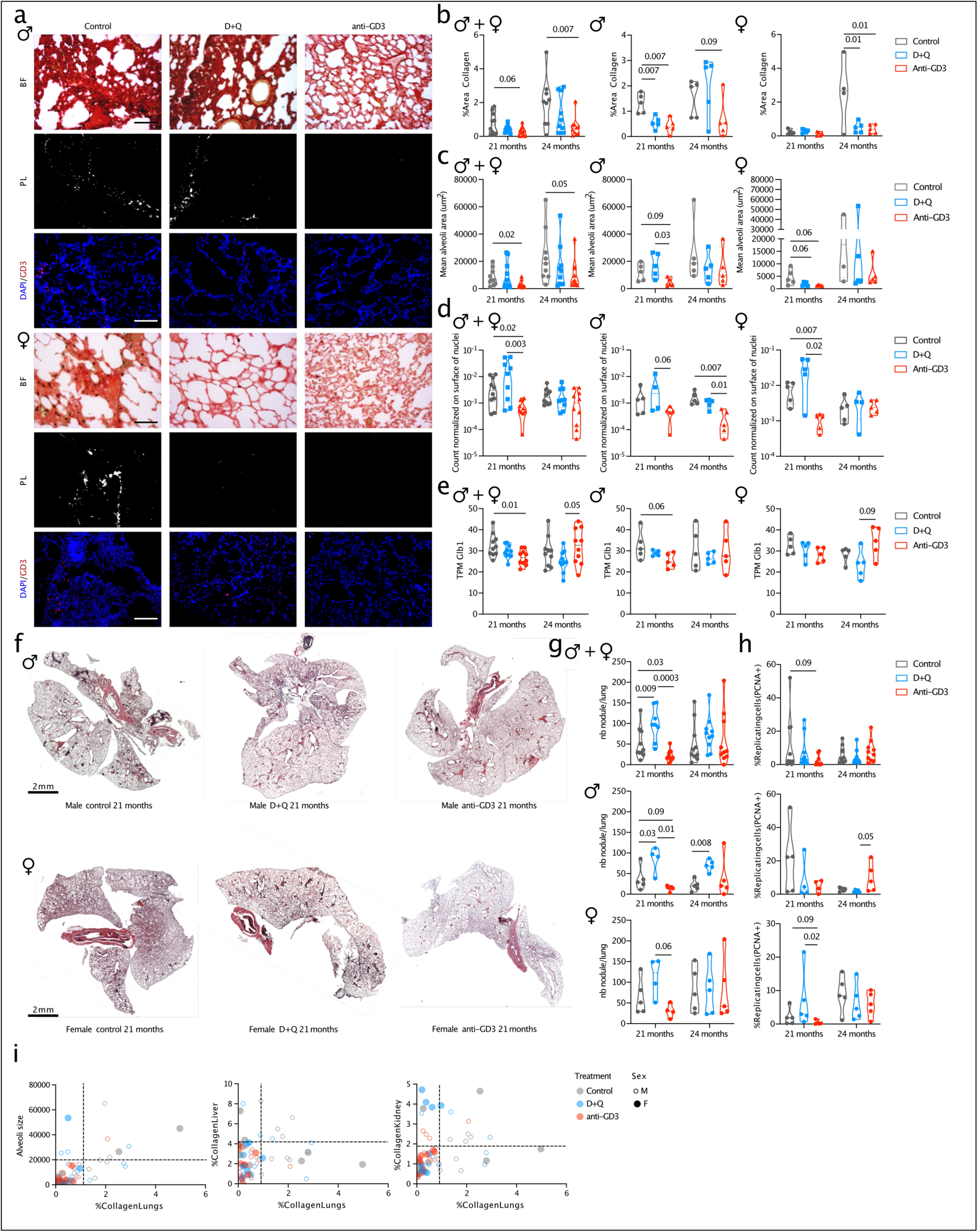
Anti-GD3 treatment blunts age-related lung pathologies in both sexes during and after cessation of treatment. **a,** Representative images of Picrosirius Red staining and GD3 immunofluorescence staining in male (upper panel) and female (lower panel) lungs at 24 months of age (3 months after treatment cessation) as a function of treatment (scale = 100 μm). Quantification of collagen percentage (**b**), alveolar size (**c**), and GD3 (**d**) in all (left panel), male (middle panel) and female (right panel) lungs as a function of treatment. **e,** Evaluation of Glb1 expression (TPKM, Transcripts Per Kilobase Million) by RNA sequencing of all (left panel), male (middle panel) and female (right panel) lungs as a function of treatment. **f**, Representative images of hematoxylin and eosin staining on male (upper panel) and female (lower panel) lungs at 21 months of age (after 3 months of treatment) as a function of treatment (scale = 2 mm). Quantification of nodules (**g**) and PCNA-positive cells (**h**) in the lungs of all (upper panel), male (middle panel) and female (lower panel) mice as a function of treatment. **i**, Correlation between collagen percentage in the lungs and alveolar size (left panel), liver collagen percentage (middle panel) and kidney collagen percentage (right panel). The experiment was performed with n=5 mice per sex and per treatment. Statistical analysis: Mann-Whitney (b-h).

We then investigated the effects of anti-GD3 in other age-associated fibrotic organs including liver and kidneys. In the liver, anti-GD3 treatment attenuated age-related fibrosis to a greater extent in male than in female mice (Extended Data Fig. 2a–c). This correlates with a sustained reduction in hepatic GD3 expression observed exclusively in males. These sex-specific responses suggest a differential regulation or clearance of GD3+ senescent cells in the liver. Interestingly, when investigating age-related kidney pathologies, we found that the kidneys of aged female mice were selectively protected by anti-GD3 therapy. Anti-GD3 treatment limited the progression of renal interstitial fibrosis exclusively in female mice (Extended Data Fig. 2d, e). However, no significant preventive effect was observed on glomerular enlargement, although there was a non-significant trend toward limiting glomerular detachment from Bowman’s capsule (Extended Data Fig. 2f, g). Despite more visible effects in female mice, we observed a decrease in GD3 expression in both sexes correlating with a reduction in renal fibrosis, consistent with reduced local senescent cell burden. (Extended Data Fig. 2a, h). Interestingly, by correlating disease burden in treated animals compared to control, we observed that the therapeutic benefit of anti-GD3 was not confined to isolated tissues. Indeed, animals that responded with reduced lung fibrosis often showed concomitant improvement in emphysema, liver, and kidney fibrosis (Fig. 3i), suggesting a systemic impact of transient GD3-targeting on senescence-driven tissue remodeling. Together, these results demonstrate that anti-GD3 therapy confers broad, organ-specific protection against age-related degeneration, with sex-dependent differences in durability and target organ sensitivity.

### Anti-GD3 treatment induces sex-specific transcriptional remodeling of the aging lung with limited overlap with senolytic pathways

To investigate the molecular mechanisms underlying the histological improvements observed with anti-GD3 therapy, we performed RNA sequencing on lung tissues from the same animals used for histopathological analysis. At 21 months of age (end of treatment), D+Q induced a strong and broad transcriptional response characterized by marked reduction of senescence-associated gene sets, including p53 and SASP signatures, and enrichment of pathways related to extracellular matrix organization and elastic fiber remodeling (Extended Data Fig. 3 and 6b). In contrast, anti-GD3 treatment induced only modest changes at this time point (Fig. 4a, b, Extended Data Fig. 5a), suggesting that its immediate mode of action does not rely on extensive gene expression reprogramming but probably relies essentially on NK cell-mediated elimination.

**Figure 4:**
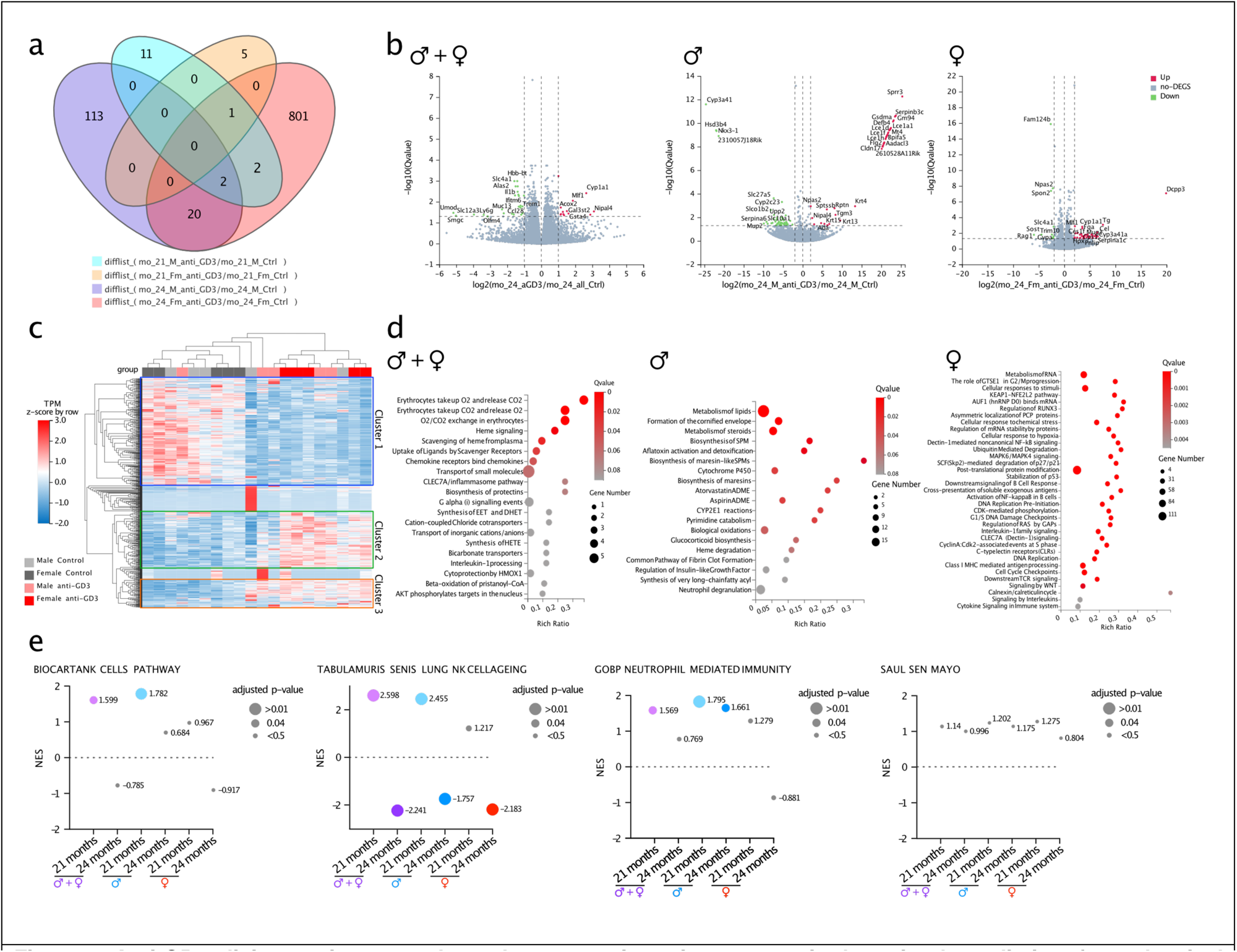
Anti-GD3 elicits a unique, sex-dependent transcriptomic response in the aging lung distinct from classical senolysis. **a**, Venn diagram of differentially expressed genes (DEG) identified by bulk RNA sequencing in anti-GD3-treated and untreated mouse lungs across both sexes at 21 and 24 months of age. **b**, Volcano plots depicting DEGs between anti-GD3-treated and untreated lungs at 24 months of age for all mice (left panel), males (middle panel), and females (right panel). **c**, Heatmap of DEGs between anti-GD3-treated and untreated mice at 24 months of age. **d**, Reactome enrichment analysis of DEGs in anti-GD3-treated lungs compared to untreated lungs at 24 months of age, shown for all mice (left panel), males (middle panel), and females (right panel). **e**, Gene set enrichment analysis (GSEA) of anti-GD3-treated compared to untreated lungs. RNA-seq was performed on lungs from n=5 mice per sex and per treatment.

However, at 24 months of age, three months after treatment cessation, anti-GD3 treatment led to significant overall and sex-specific transcriptional effects (Fig. 4a, c and d). In both sexes, treatment enhanced lung functional capacity, including signatures associated with alveolar structure maintenance and gas exchange, while selectively downregulating glycoprotein and glycolipid metabolism, most notably the ganglioside biosynthetic pathway, mirroring the sustained reduction of GD3 levels in lung tissue (Fig. 4b-e, Extended Data Fig. 5b). In male mice, anti-GD3 upregulated metabolic and mitochondrial pathways, steroid and lipid metabolism, and xenobiotic detoxification programs, together with activation of innate immune effectors such as NK cell and neutrophil-mediated immunity (Fig. 4e, Extended Data Fig. 6a). This innate immune activation was absent in females, despite increased enrichment in adaptive immune pathways, including antigen presentation, B cell activation, and immunoglobulin production (Fig. 4d, Extended Data Fig. 5c). This divergence between sexes suggests distinct immunological mechanisms of action, with males favoring cytotoxic innate surveillance and females preferentially mobilizing adaptive responses. Comparative analyses between lungs from anti-GD3 and D+Q confirmed that these treatments act through largely non-overlapping pathways. While D+Q responses were immediate, strongly senescence-focused, and associated with extracellular matrix and elastic fiber remodeling, anti-GD3 effects emerged later, targeted ganglioside metabolism, and exhibited marked sexual dimorphism in immune activation (Extended Data Fig. 4, 5c and 6c). These observations highlight distinct mechanisms of action between D+Q and anti-GD3 despite shared benefits for age-related lung pathology. Together, these findings indicate that anti-GD3 reshapes lung aging and decreases age-related lung pathologies through delayed, targeted transcriptional remodeling that aligns with its persistent histological benefits, while engaging distinct and sex-specific immune programs.

To conclude, we propose an antibody targeting the senescence-associated ganglioside GD3 as one of the first immunotherapies to prevent pathological aging. Unlike other senolytic treatments that require chronic administration^4,14,15^, a short mid-life treatment conferred durable benefits for over a year, attenuating pulmonary, hepatic, and renal pathologies, and improving both lifespan and healthspan. The effects were mechanistically distinct from D+Q^16^ and showed marked sexual dimorphism, with males primarily protected from non-cancer mortality via metabolic and innate immune activation, and females from tumor-related mortality through adaptive immune engagement. These findings establish GD3 as a therapeutic senescence-associated immune checkpoint in aging and underscore the importance of identifying additional senescence-induced immune checkpoints to develop next-generation senescence-targeted immunotherapies and highlight the critical importance of considering sexual dimorphism in geroscience research.

## Acknowledgments

We extend our gratitude to the IRCAN core facilities (animal facility and PICMI) made possible through funding from FEDER, the Ministry of Higher Education, the Provence-Alpes-Côte d’Azur Region, the Conseil Départemental 06, Cancéropole PACA, the French National Cancer Institute (INCA), and the Association pour la Recherche sur le Cancer (Fondation ARC). We would like to express our warmest thanks to Thierry Juhel for his invaluable assistance in monitoring, treating and analyzing mouse experiments. We thank Valérie Duranton-Tanneur for her help with RNA extraction from FFPE samples. This work was conducted through the Adipo-Cible Research Study Group, supported by the French government through the France 2030 investment plan managed by the National Research Agency (ANR), as part of the Initiative of Excellence of Université Côte d’Azur under reference number ANR-15-IDEX-01 and the Institut Hospitalo Universitaire IHU Respirera supported by the French National Research Agency (ANR), ANR-23-IAHU-007.

## Funding

Longevity Impetus Grant (contract # 248181) (JCV)

Cancéropôle PACA, Région Sud Provence-Alpes-Côte d’Azur, and INCa N°2024-16Kpole (JCV)

French National Cancer Institute (INCA) AAP PLBIO 2023 (PLBIO23-100-2023-159) (JCV)

French National Research Agency, (ANR) PRC SENEDIT ANR-22-CE13-0011-01 (JCV)

French National Research Agency, (ANR) PRC FibroTargLung ANR-23-CE19-0038-04 (JCV)

French National Research Agency, (ANR), France 2030 ANR-23-IAHU-0007 (JCV)

French Ministry of Research (MNRT) PhD Fellowship 2021-2024 (IM)

Fondation ARC pour la recherche sur le cancer; PhD Fellowship 2024-2025 ARCDOC42024020007920 (IM)

## Author contributions

Conceptualization: J.C-V

Data curation: J.C-V, I.M, L.O., L.B., B. DM.

Formal analysis: J.C-V, I.M, L.O., L.B., B. DM.

Funding acquisition: J.C-V

Investigation: I.M, T.J, L.O., R.R, L.B., B. DM, J.C-V.

Methodology J.C-V, I.M, L.O.

Project administration: J.C-V

Resources: E.G, J.C-V

Software: L.O, G.C

Supervision: J.C-V

Validation: J.C-V, I.M, L.O., G.C, L.B., B. DM

Visualization: I.M, J.C-V

Writing – original draft: I.M, J.C-V

Writing – review & editing: G.C, I.M, E.G, L.B., B. DM and J.C-V.

---

**Extended Fig. 1:**
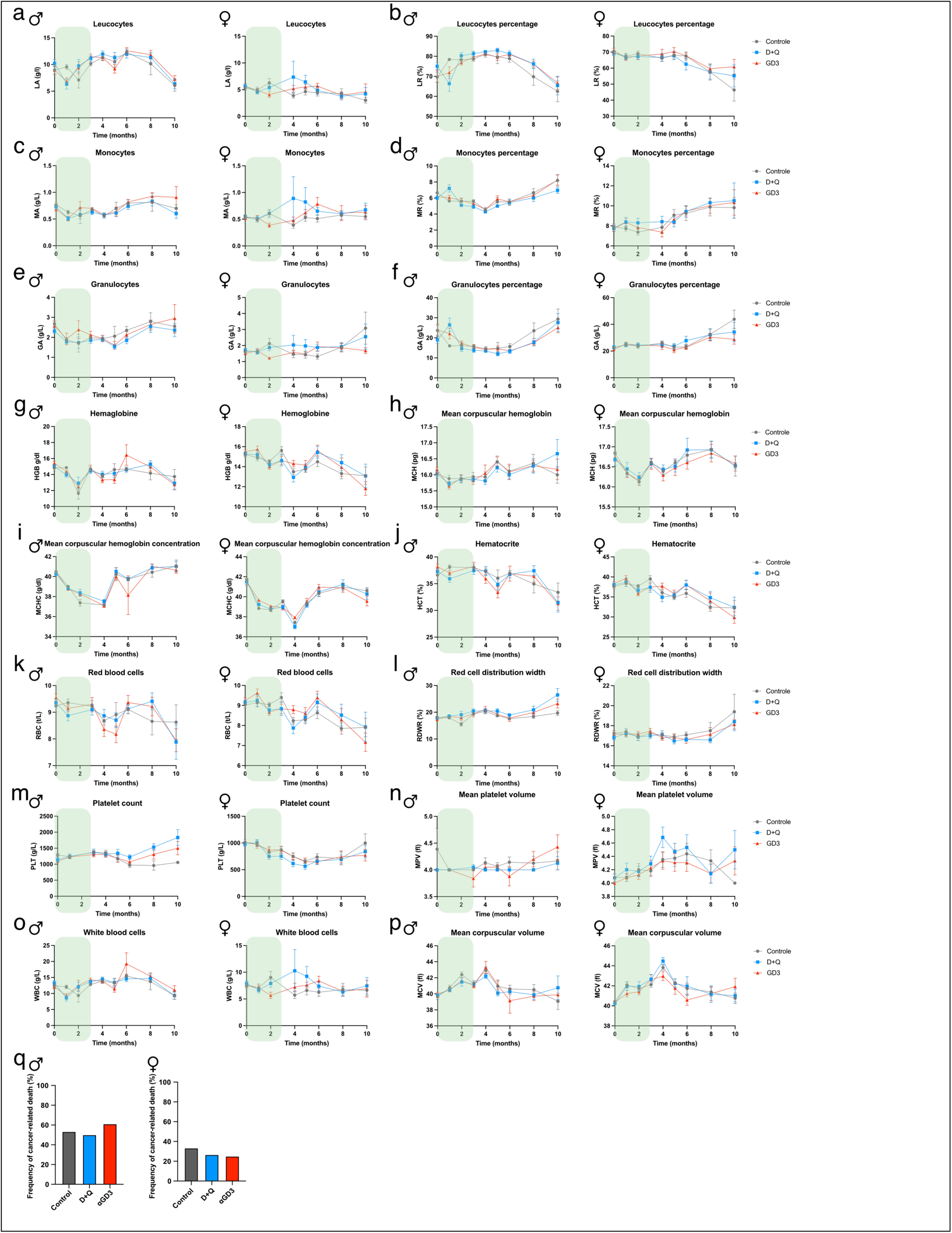
Anti-GD3 treatment does not affect hematological parameters in male and female mice. **a-f**, Quantification of absolute (LA, **a**) and relative (LR, **b**) leucocyte counts, leucocytesabsolute (MA, **c**) and relative monocyte counts (MR, **d**), and absolute (GA, **e**) and relative granulocyte counts (GR, **f**) in blood. The green shaded area indicates the treatment period. **g-p**, Analysis of hematological parameters: hemoglobin (HGB, **g**), mean corpuscular hemoglobin (MCH, **h**), mean corpuscular hemoglobin concentration (MCHC, **i**), hematocrit (HCT, **j**), red blood cells (RBC, **k**), red cell distribution width (RDW, **l**), platelet count (PLT, **m**), mean platelet volume (MPV, **n**), white blood cells (WBC, **o**) and mean corpuscular volume (MCV, **p**).. **q**, Percentage of cancer-related death stratified by treatment, in male (left) and female (on the right) mice. For all panels, n=25 mice per sex per treatment group. Data are presented as mean ± s.e.m. or s.d (panel a-p). Statistical tests: Two-Way ANOVA (a-p), Mann-Whitney U-tests (q).

**Extended Fig. 2:**
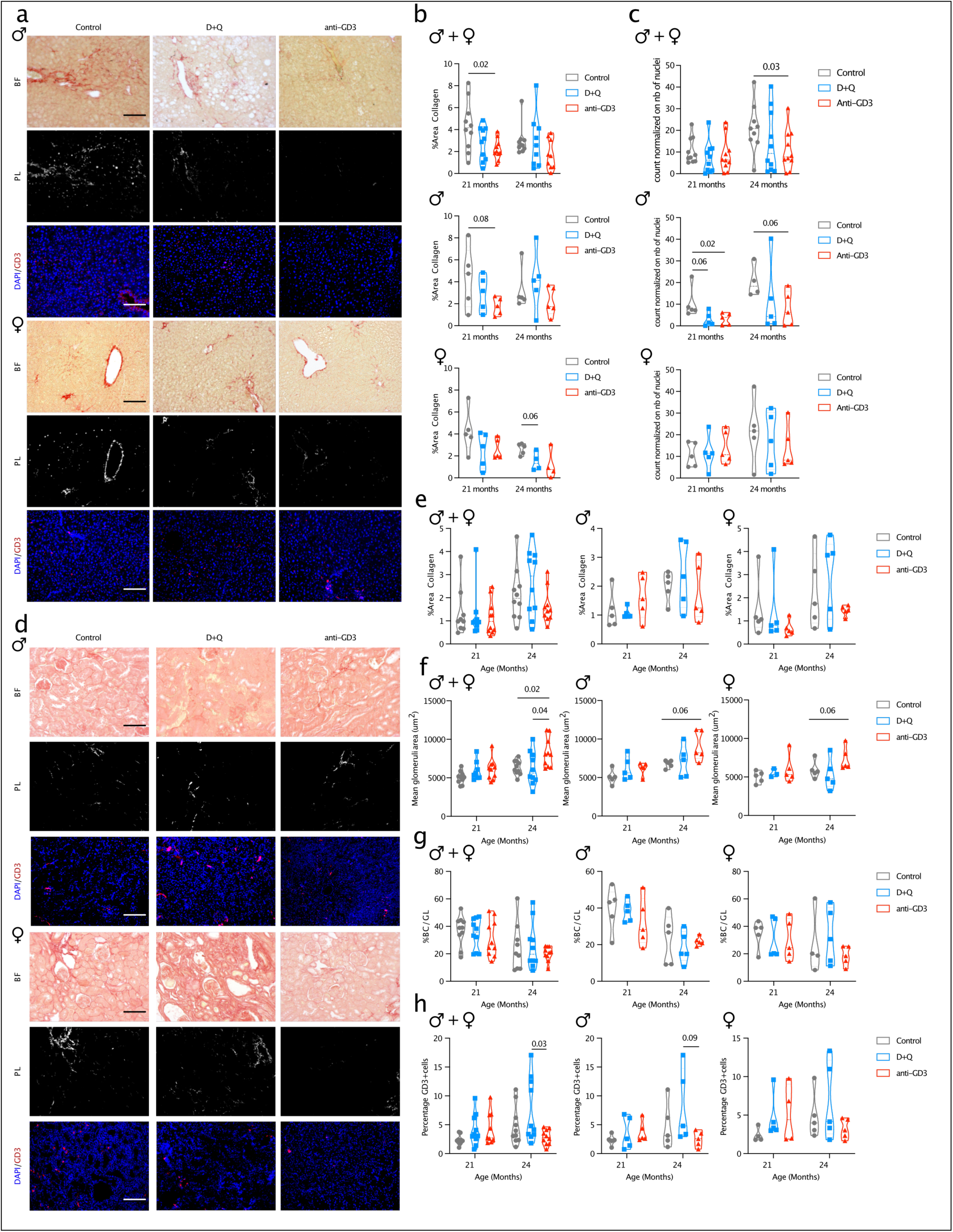
Anti-GD3 treatment blunts the development of age-related liver and kidney fibrosis in a sex-dependent manner. **a**, Representative staining for collagen (Picrosirius Red, detected by polarized light (PL)) and GD3 (immunofluorescence) in liver sections from 24-month-old male (top) and female (bottom) mice, 3 months after the end of the treatment. **b, c**, Quantification of the percentage of collagen-positive area (**b**) and GD3 signal (**c**) in the liver, analyzed for the combined cohort (top) and separately for males (middle) and females (bottom). **d**, Representative staining for collagen (Picrosirius Red, detected by polarized light (PL)) and GD3 (immunofluorescence) in kidney sections from 24-month-old male (top) and female (bottom) mice, 3 months after the end of the treatment. **e, h**, Quantification of renal fibrosis and morphology, including collagen-positive area (**e**), glomeruli size (**f**), the Bowman’s capsule to glomeruli area ratio (**g**), and GD3 signal (**h**), analyzed for the combined cohort (left) and separately for males (middle) and females (right). Experiments were performed with n=5 mice per sex per treatment group. Statistical analysis: Mann-Whitney (b-h). All scale bars represent 100μm. BF: bright field; PL: polarized light.

**Extended Fig. 3:**
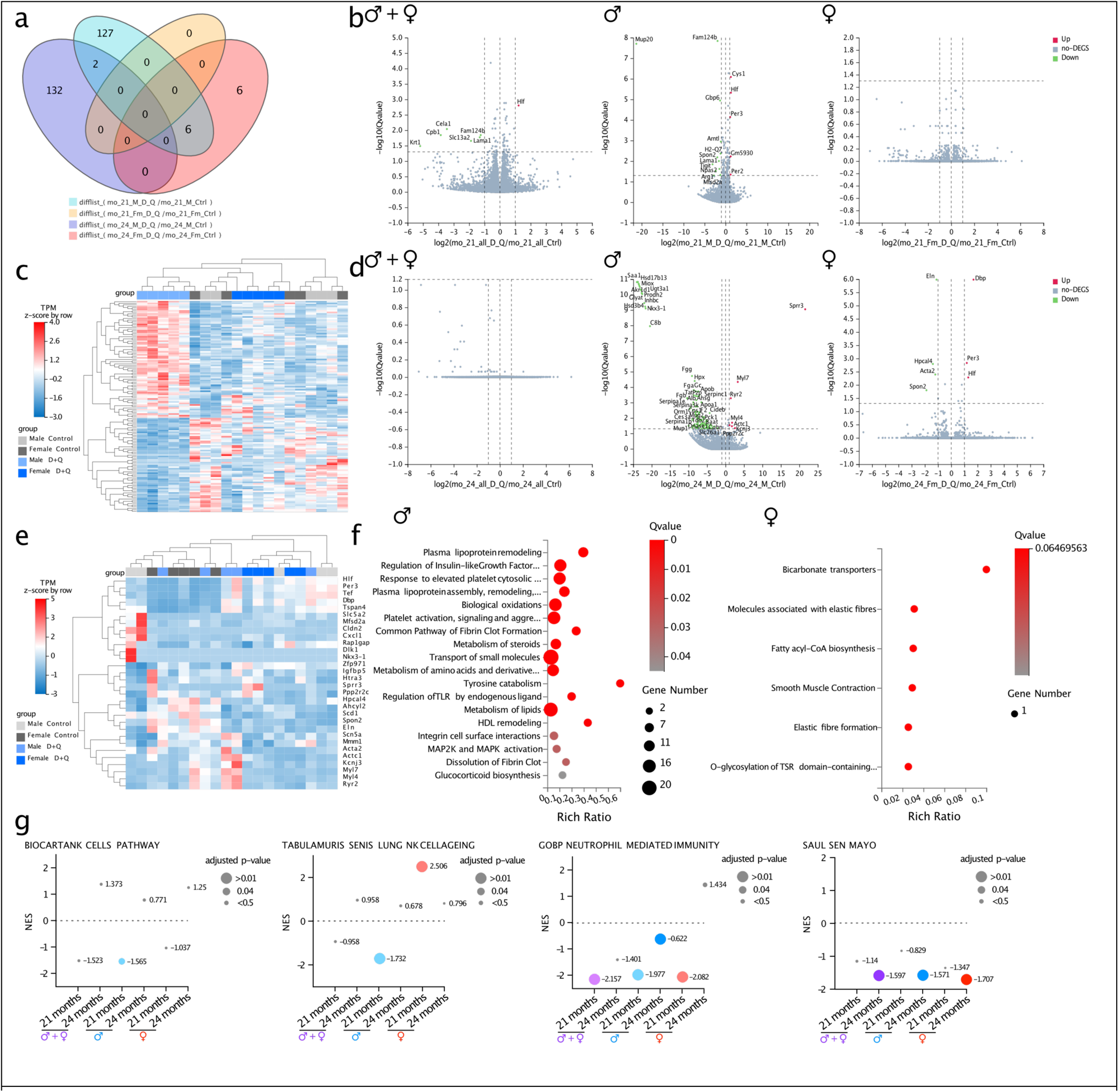
Sex-specific transcriptomic response to D+Q senolytic therapy in aged lungs. **a**, Venn diagram showing the overlap of differentially expressed genes (DEGs) identified by bulk RNA-seq of lungs between D+Q-treated and untreated mice of both sexes at 21 and 24 months of age. **b-e**, Volcano plots (**b, d**) and heatmaps (**c, e**) of DEGs in lung tissue between D+Q-treated and untreated mice at 21 months (**b, c**) and 24 months (**d, e**). Analyses are shown for the combined cohort (left), males only (middle), and females only (right). **f**, Reactome pathway enrichment analysis of DEGs in lung tissue in D+Q-treated vs untreated animals at 24 months, shown for males (left) and females (right). **g**, Gene set enrichment analysis (GSEA) of lung tissue from D+Q–treated versus untreated mice, stratified by sex and timepoint. All analyses were based on bulk RNA-seq of lung tissue from n=5 mice per sex and per treatment group.

**Extended Fig. 4:**
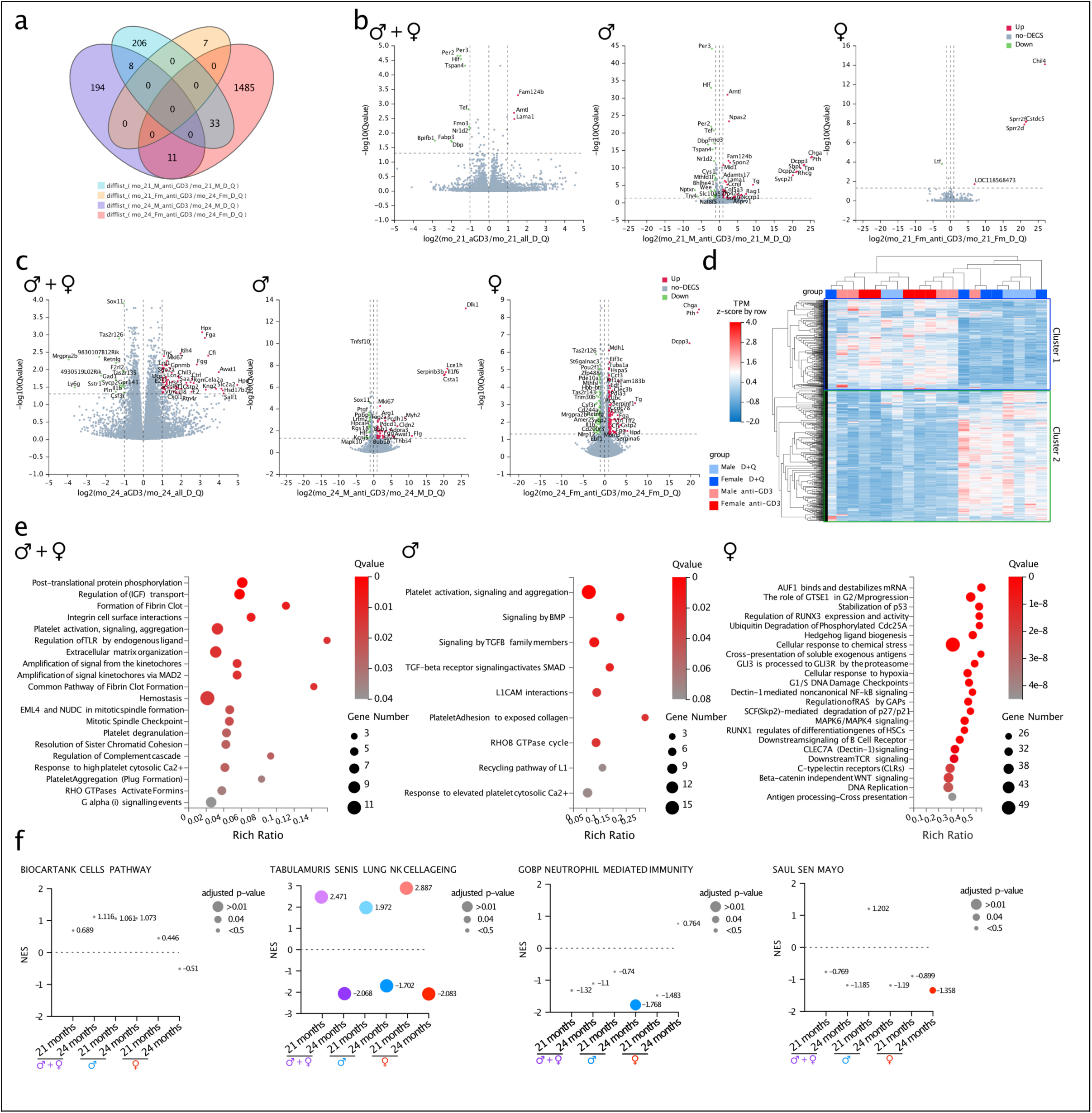
Anti-GD3 and D+Q treatments elicit distinct, sex-specific transcriptomics signatures in the aged lung. **a**, Venn diagram showing the overlap of differentially expressed genes (DEGs) identified by bulk RNA-seq of lungs between anti-GD3- and D+Q-treated mice of both sexes at 21 and 24 months of age. **b,c**, Volcano plots of DEGs in lung tissues between anti-GD3- and D+Q-treated mice at 21 months (**b**) and 24 months (**c**) of age. Analyses are shown for the combined cohort (left), males only (middle) and females only (right). **d**, Heatmap of DEGs in lung tissue between anti-GD3- and D+Q-treated mice at 24 months of age. **e**, Reactome pathway enrichment analysis of DEGs in lung tissue in anti-GD3-treated vs D+Q-treated mice at 24 months of age. Analyses are shown for the combined cohort (left), males only (middle) and females only (right). **f**, Gene set enrichment analysis (GSEA) of lung tissue from anti-GD3-treated versus D+Q–treated mice, stratified by sex and timepoint. All analyses were based on bulk RNA-seq of lung tissue from n=5 mice per sex and per treatment group.

**Extended Fig. 5:**
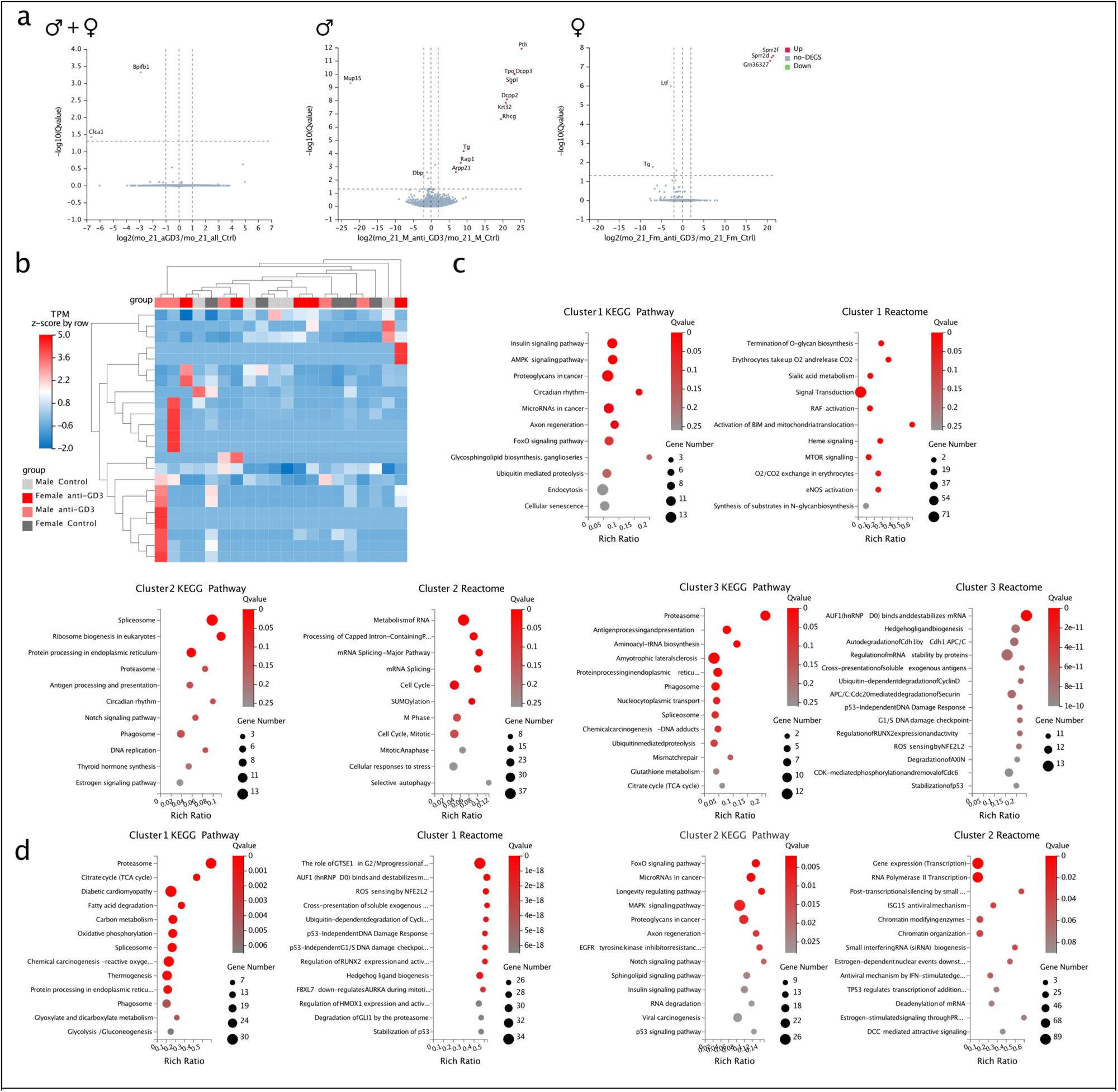
Anti-GD3 treatment impact ganglioside metabolism, immune, and cell cycle pathways in the aged lung. **a,** Volcano plots depicting DEGs in lung between anti-GD3-treated and untreated mice at 21 months of age. b, Heatmap of DEGs between anti-GD3-treated and untreated mice at 24 months of age. c, d, KEGG pathway and Reactome enrichment analyses for gene clusters modulated by anti-GD3 treatment relative to untreated controls (**b,** Fig. 3d) and relative to D+Q treatment (**c,** Ext. Fig. 4d) at 24 months. All analyses were based on bulk RNA-seq of lung tissues from n=5 mice per sex and per treatment group.

**Extended Fig. 6:**
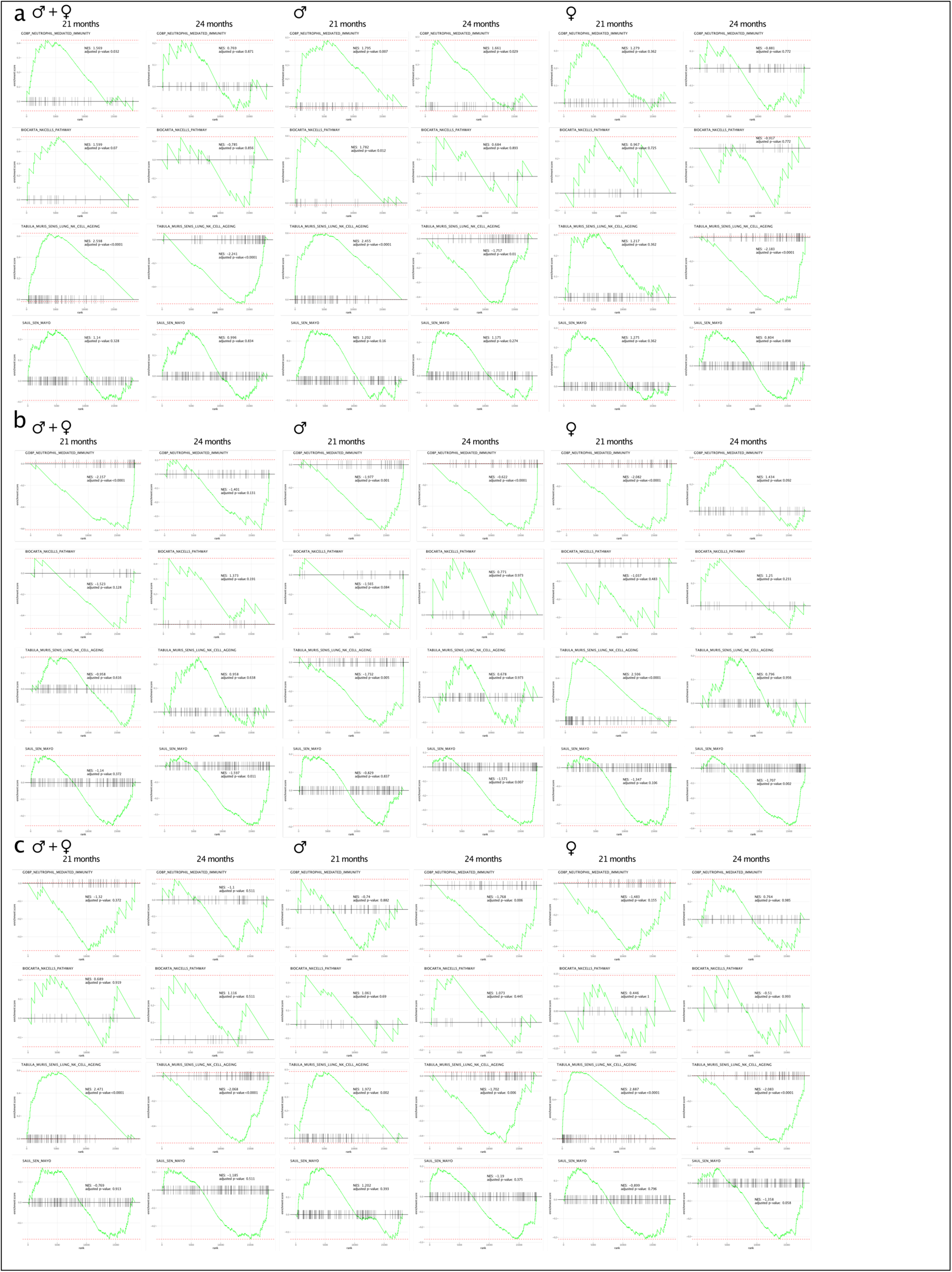
Anti-GD3 and D+Q treatments differentially regulate immune and senescence gene sets. **a-c**, Gene Set Enrichment Analysis (GSEA) plots for four key pathways: GOBP NEUTROPHILE MEDIATED IMMUNITY, BIOCARTA NKCELLS PATHWAY, TABULA MURIS SENIS LUNG NK CELL AGEING and SAUL SEN MAYO (cellular senescence). The plots show enrichment in three comparisons: anti-GD3 treated vs untreated (**a**), D+Q-treated vs untreated (**b**), and anti-GD3-treated vs D+Q-treated (**c**). For each comparison, plots are shown for the combined cohort (left), males only (middle) and females only (right). Within each of these groups, data are presented for both 21 (left) and 24 (right) months of age. These plots provide detailed views of data summarized in Fig. 3f, Ext. Data Fig. 3g and Ext. Data4f. All analyses were based on bulk RNA-seq of lung tissue from n=5 mice per sex and per treatment group.

**Extended Data 1: Frailty index scoring parameters.**

**Extended Data 2: Differential Gene Expression between different treatment groups, sexes and time points.**

## Material and Methods

The research conducted in this study complies with local and institutional guidelines. The study protocol, including all procedures involving animals, was reviewed and approved by the "Comité Institutionnel d’Éthique Pour l’Animal de Laboratoire," registered at the French Ministry of Higher Education and Research under N° 28. The research was conducted under authorization n° MESR 2022012015452005.

### Treatment of 18-month-old mice

Animals were maintained on a 12∶12-h light-dark cycle with food and water provided ad libitum. Pathogen-free 18-month-old male and female C57BL/6J mice (Charles River) were treated six times, every 15 days, with 150 µg of anti-GD3 (BIOTEM production, clone R24) in 150 µl of phosphate-buffered saline (PBS) via intraperitoneal injection (I.P.) and 500 µl of sterile water by oral gavage or with Dasatinib (Bio-Techne ref# 6793/100) (5 mg/kg) and Quercetin (Bio-Techne ref# 1125/100) (50 mg/kg) by oral gavage in 500 µl of sterile water and with 150 µg of isotype IgG3 (BIOTEM production) in 150 µl of phosphate-buffered saline (PBS) via intraperitoneal injection (I.P.). For all mouse experiments, the number of animals required was determined using a Monte Carlo power test prior to the experiments. All mouse experiments were conducted in compliance with local and international institutional guidelines and were approved by the Animal Care Committee of IRCAN, as well as regional (CIEPAL Côte d’Azur Agreements NCE/2015-266#2015102215087555 and NCE/2020-675#2020042723583497) and national (French Ministry of Research) authorities.

### Lifespan study

Mice were randomized into treatment groups at the cage level to ensure comparable initial weights and frailty indices across all groups. Euthanasia was performed for animals that became moribund or exhibited predefined health concerns, such as significant tumor burden, paralysis, or rapid weight loss. Daily inspections were conducted, and deceased mice were recorded and stored in a refrigerator for necropsy. During necropsy, tissues were examined for tumors, spleen enlargement, and infections, with cancer presence or absence documented accordingly. A "cause of death" was assigned to each necropsied mouse based on the most prominent findings observed.

### Tissue collection

Tissues were collected 3 months after the initiation of the treatment and 3 months after treatment cessation. Lung, kidney, and liver tissues were excised and processed for either cryopreservation (included in OCT freezing medium) or histological analysis (fixed with 4% (w/v) formaldehyde in PBS, embedded in paraffin, sectioned, and stained with hematoxylin/eosin and picrosirius red).

### Histology & immunofluorescence

For murine tissues, antigen retrieval was performed on 5-μm paraffin sections using a Vector unmasking reagent (Vector Laboratories, ref# H3300). Sections were blocked (Avidin/Biotin blocking kit, MOM kit, Vector Laboratories, ref# BMK-2202 and SP-2001) and incubated overnight at 4°C with mouse monoclonal anti-GD3 (Abcam, ref#ab11779, 1:40) or rabbit anti-PCNA (Invitrogen, ref#13-3900, 1:100). Detection of primary antibodies involved the use of a biotinylated anti-mouse IgG (MOM kit) or anti-rabbit IgG (Jackson ImmunoResearch, ref# 111-065-144). For immunofluorescence, tissues were incubated with streptavidin-Cy3 (Jackson ImmunoResearch, ref# 016-160-084) for 1 hour at room temperature. Stained tissue sections were sequentially scanned using an HD Zeiss microscope, enabling imaging of entire sections. The percentage of collagen was quantified using ImageJ, while tumor numbers and immunofluorescence data were quantified using QuPath.

### Bulk RNA-seq on FFPE lung samples

RNA extraction followed the Maxwell RSC RNA FFPE kit (Promega, ref# AS1440) protocol, with quality and concentration assessed via Nanodrop and Bioanalyzer (Agilent). Library preparation, sequencing, and initial data filtering, including adaptor removal, were conducted by BGI Genomics. Sequencing was performed on the DNBSEQ platform in paired-end mode with 150 bp reads. Low-quality reads, adaptor contamination, and excessive unknown bases were filtered out using SOAPnuke. HISAT2 was used to align reads to the Mus_musculus_NCBI_GCF_000001635.27_GRCm39 genome, followed by fusion gene and differential splicing detection with Ericscript (0.5.5-5) and rMATS (v4.1.1). Further alignment to the gene set was performed with Bowtie2, and quantification was done using RSEM (v1.2.28). Differential gene expression analysis was conducted with DESeq2. Downstream analysis, including differential expression and heatmap generation, was performed using BGI’s Dr. Tom software, with heatmaps generated from log-normalized TPM values. Gene-set enrichment analysis was conducted using the fgsea package for three senescence-related pathways (SAUL_SEN_MAYO, FRIDMAN_SENESCENCE_UP, and REACTOME_SENESCENCE_ASSOCIATED_SECRETORY_PHENOTYPE_SASP) from MSigDB, considering pathways significantly enriched at an adjusted p-value (p adj) < 0.05.

### Statistics & reproducibility

Sample sizes were not predetermined by a statistical method, but were chosen on sample sizes used in similar previous studies. Mouse experiments were performed on n = 25 animals per group for lifespan studies and n = 5 animals per group for histological and transcriptomic analyses, as indicated in the relevant figure legends. No exclusion criteria were defined, and no data were excluded from the analyses. Investigators were not blinded to group allocation during experiments and outcome assessment. Statistical tests were all performed using GraphPad Prism versions 10 including the normality and lognormality tests, the Mann-Whitney U-tests, the log-rank tests, Pearson correlations, and two-way ANOVA tests.

### Use of Large Language Models

ChatGPT-4o (OpenAI) was not used for *de novo* text generation. It was used to correct the spelling, grammar and syntax of the manuscript.

